# Pervasive Intraspecific Genetic Local Adaptation Within the Natural Gut Microbiome

**DOI:** 10.1101/2025.06.06.658271

**Authors:** Russ J. Jasper, Adamandia Kapopoulou, Stephan Peischl, Bahtiyar Yilmaz, Jens Becker, Claudia Bank

## Abstract

Gut microbial communities are shaped by the physicochemical gradients and compartmentalization along the gastrointestinal tract. Whether spatially variable selection is strong enough to promote lasting genetic divergence within microbial species subject to migration and clonal expansion remains unclear. We performed a metagenomic population-genetic analysis of roe deer microbiota across metabolically and structurally distinct gut compartments. Remarkably, 10 of 20 prevalent bacterial species exhibited clear signatures of genetic local adaptation, despite the homogenizing effects of frequent migration. Nutrient availability and environmental stability emerged as primary drivers of selection, with horizontal gene transfer as a key mechanism enabling local adaptation. Our findings demonstrate that a comprehensive understanding of ecological and evolutionary dynamics in the gut necessitates considering fine-scale genetic variation within species.

---

The gut microbiome plays a central role in host physiology, development, and health, and is increasingly recognized as an evolutionary ecosystem shaped by both ecological and genetic forces (1). Understanding how microbial populations evolve within the spatially heterogeneous environment of the gastrointestinal tract remains a fundamental challenge in microbiome biology and host-associated population genetics. Although distinct gut compartments present divergent physicochemical conditions that could impose localized selective pressures, it remains unclear whether such heterogeneity drives within-species genetic divergence. Do gut microbes undergo genetic local adaptation, whereby allelic variants become associated with specific environmental niches, or do homogenizing forces such as strong migration and clonal reproduction erode any possible regional differentiation? Establishing whether genetic divergence within microbial species reflects adaptation to the compartmentalized environments of the gut is critical for linking microbial genomic variation to ecological function. Most prior research on microbiome responses to spatial heterogeneity in the gut has focused on species-level or higher taxonomic resolution rather than intraspecific genetic variation; however, such studies often indicate that *community-level* local adaptation occurs. Significant differences in communities between the upper and lower intestinal tract in various species (2–5) point to strong divergent selection across gut regions, raising the possibility that similar spatially structured selective pressures could drive genetic local adaptation within individual microbial species. However, comparisons between adjacent regions, such as the cecum and colon (4), proximal and distal colon (2), or colon and rectum (5), often reveal minimal community differentiation, despite distinct physiological functions and abiotic conditions. Although these patterns offer indirect evidence of ecological sorting (or lack thereof), they do not address whether genetic local adaptation occurs within species, underscoring the need for population-genomic resolution.

To date, relatively few studies have examined the adaptation of microbial species to heterogeneous gut environments at the genetic level. Recent work investigating microbiota from the human mid and hindgut focused on whether environmentally associated strains arose from single or multiple colonization events (6). The researchers found no environment-associated gains or losses in accessory genes, suggesting an absence of genetic local adaptation. However, as this was not the primary focus of their study, they did not assess local adaptation at finer genomic scales, such as single nucleotide polymorphisms (SNPs). Further work, in a lab setting, explored how human microbiota inoculated into germ-free mice adapted along the mid and hindgut (7). Here, the authors found very few SNPs that were associated with the environment and none that were consistently associated across multiple replicates. Moreover, the authors reported that community membership and abundance were associated with the environment, but that within-species genetic variation was generally unchanged regardless of environment. However, because this study explored human microbiota inoculated into mice hosts, it is also possible that any potential signal of local adaptation to the gut environment was swamped out by host-specific adaptation. Interestingly, both studies concluded that multiple, genetically distinct strains can coexist across multiple environments without spatial segregation, perhaps suggesting that genetic local adaptation is unlikely to occur in gut microbiota species (6,7).

In contrast to the above studies and taking a longitudinal sampling approach, controlled long-term experiments in gnotobiotic mice revealed signatures of positive selection and sub-strain diversification within a stable gut microbiota in the cecum (8), perhaps indicating that local adaptation is possible under reduced ecological noise, underscoring the need to test whether such processes occur in natural microbiomes shaped by more complex host and environmental heterogeneity. Additional research at the genetic level and without spatial variation has revealed substantial signals of positive or purifying selection, both within host through time (9,10) and between hosts (9–11) in human microbiota. Taken together, these studies highlight the rapid pace and the critical role of global adaptation during the evolution of microbial populations but provide no conclusive answer to whether there is relevant local adaptation of microbial populations within the intestinal tract. To the best of our knowledge, no studies have yet directly investigated whether genetic local adaptation at SNP resolution occurs in natural gut microbiome populations.

To explore how natural microbiota populations adapt to the environmental heterogeneity along the gut, we sampled microbiome communities from ten different locations longitudinally along the gastrointestinal tracts (Fig. 1A) of seven roe deer (*Capreolus capreolus*) near Biel, Switzerland on November 21, 2022. We performed metagenomic sequencing and recovered high-confidence near-complete genomes (i.e., mean completeness > 85% and mean contamination < 2%) from a total of 909 unique species (Fig. S1). Broadly, we observed three significantly different microbial community regimes along the gastrointestinal tract corresponding to the foregut, midgut, and hindgut compartments (Fig. 1B-C; Fig. S2; p<0.001), consistent with findings from previous ruminant studies (5). Next, we investigated whether local adaptation occurred within species at the genetic level. We focused our analyses on microbial populations from the hindgut, specifically the cecum and rectum, based on two main considerations. First, microbial community composition across the hindgut was homogeneous at both the taxonomic and functional levels, minimizing the confounding effects of compositional turnover and allowing us to attribute any observed genetic divergence more confidently to environmental selection rather than community restructuring. Second, the cecum and rectum represent structurally and functionally conserved compartments across mammalian hosts, including both ruminants and monogastrics. Unlike the foregut of ruminants, which is highly specialized for pregastric fermentation and exhibits substantial anatomical and microbial divergence, the hindgut is more evolutionarily and physiologically comparable across taxa. By targeting these two homologous compartments, our findings are more likely to be generalizable beyond ruminants and offer broader insights into the evolutionary dynamics of gut microbiota in spatially structured environments.

**Figure 1.**
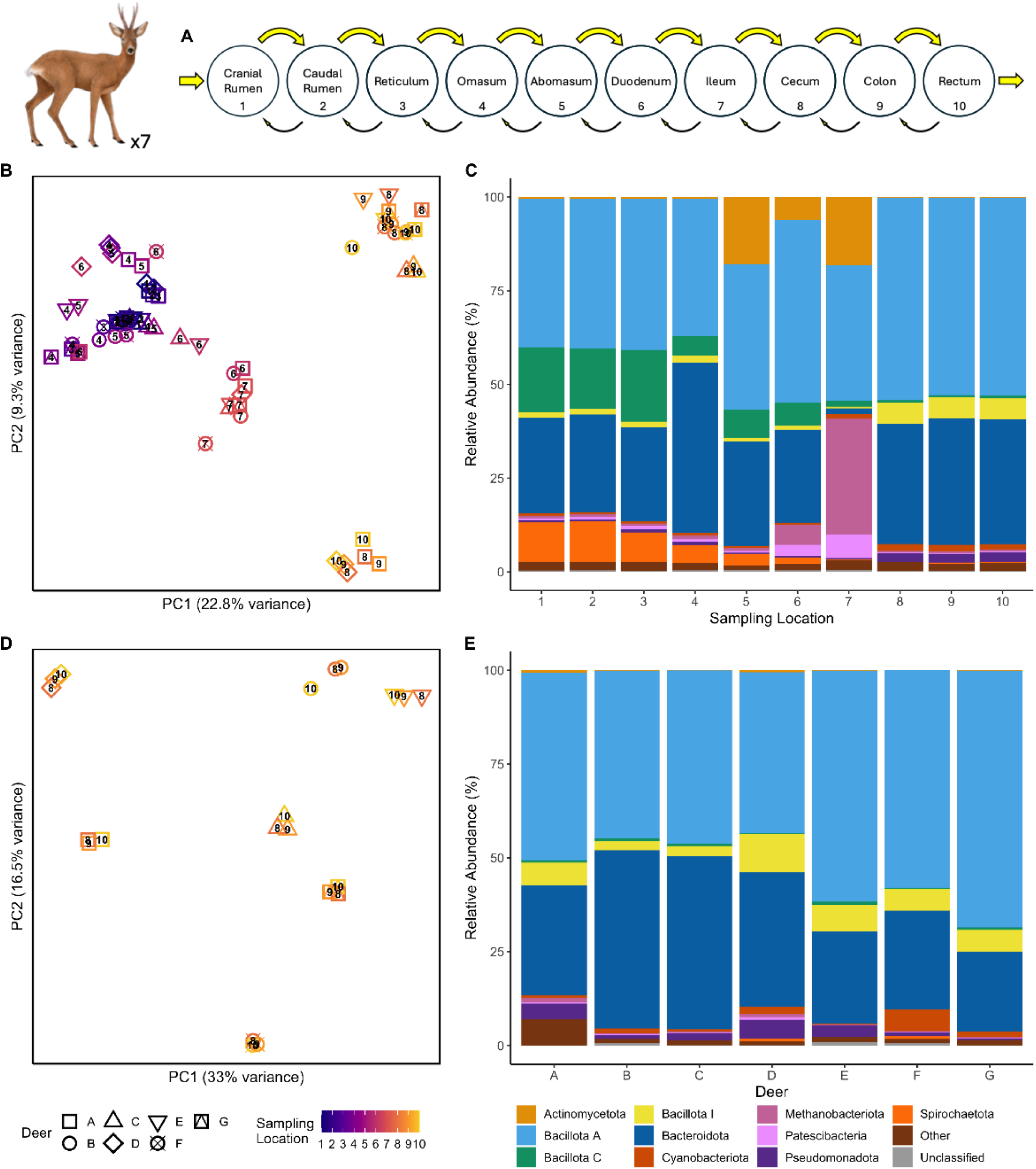
Microbial community structure along the gastrointestinal tract of seven wild deer reflects environmental and host-specific factors. (A) Schematic of the roe deer gastrointestinal tract showing all 10 sampling locations; arrows indicate hypothesized migration rates between gut compartments. (B-C) Along the gastrointestinal tract, community composition is primarily structured by compartment: principal component analysis (PCA) of species-level relative abundances (B) and corresponding phylum-level composition profiles grouped by gut region (C). (D-E) In contrast, within the hindgut (locations 8-10), composition is primarily structured by host: PCA of species-level relative abundances (D) and corresponding phylum-level composition profiles grouped by individual deer (E). Panels C and E show the 10 most abundant phyla with the remaining taxa grouped as ‘Other’. The ‘Other’ category was composed of Verrucomicrobiota, Elusimicrobiota, Synergistota, Campylobacterota, Halobacteriota, Bacillota, Myxococcota, and Desulfobacterota (ordered by overall abundance). Roe deer image credit: Adobe Stock.

We called SNPs in all hindgut populations and prioritized the top 20 species according to the number of filtered SNPs. We then conducted a stratified contingency analysis at each locus to identify SNPs that displayed signals of putative local adaptation, that is, sites with substantial allele frequency changes between cecum and rectum populations in a concordant direction across replicate deer. After correcting for false discovery (FDR threshold=0.05), we classified all SNPs that were significant and in the top 0.01% most extreme SNPs within a species as candidates for local adaptation. Strikingly, our findings revealed that 10 out of 20 microbial species exhibited significant signals of putative genetic local adaptation (Fig. 2), even in light of the remarkably homogeneous communities and across a relatively small spatial scale. This high rate of genetic local adaptation suggests that gut microbiome communities cannot be thought of as an assemblage of the same generalist genotypes distributed throughout the gut; rather, many microbial species are genetic specialists with fine-grained genetic adaptations to their local environments. The frequent occurrence of genetic local adaptation between such homogeneous communities challenges the notion that microbiomes are shaped primarily through ecological sorting. Instead, we show that local evolutionary processes played a dominant role in shaping the hindgut communities we studied. Our findings stand in stark contrast to previous microbiome research which reported little genetic differentiation across gut compartments (6,7). Finally, given that most gut microbiota species reproduce clonally and are generally considered to experience high rates of migration (e.g., (7,9,12)) our findings are perhaps unexpected and speak to the power of natural selection in shaping microbial populations.

**Figure 2:**
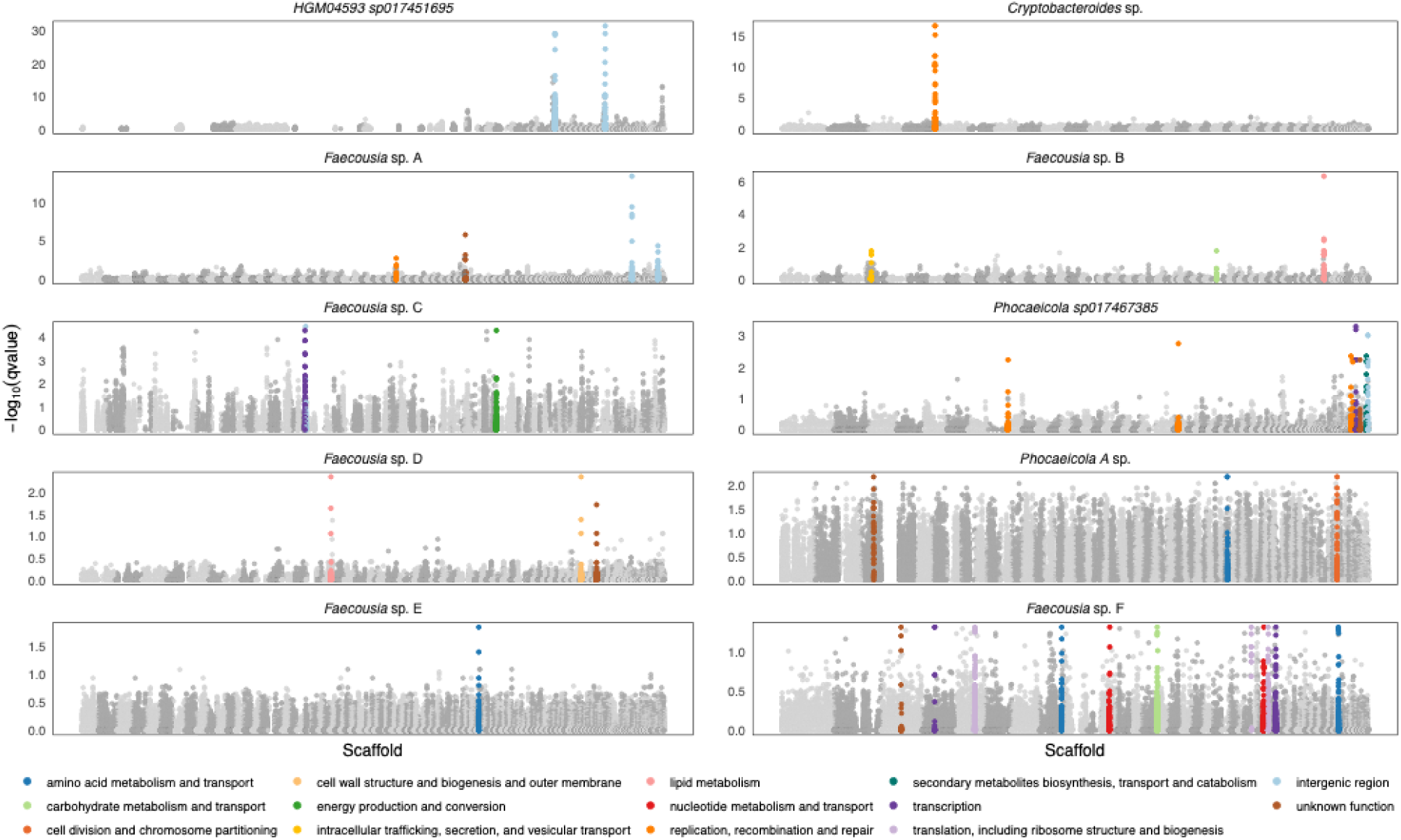
Signatures of putative genetic local adaptation are prevalent in multiple gut microbiota species. The Manhattan plots show the Cochran-Mantel-Haenszel q-value for each SNP on the y axis, comparing the cecum and rectum populations of a given microbial species across replicate deer. Each genome is at scaffold resolution; scaffolds are shown in greyscale and ordered by decreasing size. Outlier genes or intergenic windows are shown in color, with all SNPs within a given outlier region highlighted. For genic outliers, the color scale denotes their putative function according to the COG database (13). Outlier genes or intergenic windows were identified as those with significant SNP q-values (FDR=0.05) and in the top 0.01% of all SNPs for a given species. Panels are organized according to the most extreme SNP in each species. Note that the magnitude of the y-axis changes between species.

Next, we aimed to elucidate the biological processes driving the divergent selection between the cecum and rectum populations that likely gave rise to the local adaptation outliers. Among the 10 species with outlier SNPs, we inferred the functions of all genes (n=34) and intergenic regions (n=9) with outliers based on the closest known orthologous genes or nearby regulatory motifs (Fig. 2; Table S1). Among genic outliers the most frequent outlier functions were macromolecule transport and metabolism; or replication, recombination and repair (Fig. S3; COG database (13)). Intergenic outliers were associated with macro- and micronutrient metabolism; energy production; replication; and dormancy functions (14–19). Broadly, our findings are consistent with two main divergent selection pressures driving local adaptation in the deer gut: nutrients (e.g., macromolecule transport and metabolism) or adaptability (e.g., replication, recombination and repair).

The cecum and rectum represent contrasting nutrient environments: the cecum, positioned at the entrance to the colon, receives nutrient-rich input from the small intestine and serves as a major site of hindgut fermentation and microbial nutrient synthesis (20). In contrast, the rectum, located at the distal end of the colon, receives nutrient-depleted digesta and primarily functions in waste excretion, with minimal microbial fermentation and nutrient production. Our results indicate that heterogeneity in nutrients likely drove genetic local adaptation outliers in seven of the analyzed species: *HGM04593 sp017451695*; *Phocaeicola A* sp.; *Faecousia* B., C., D., E., and F. (Fig. 2; Table S1). In *HGM04593 sp017451695*, for example, we identified two compelling outlier peaks (Fig. 2, top left) with multiple outlier SNPs within or in very close proximity to sequences that significantly matched known regulatory motifs (p<0.001) Fur, PvdS, and DosR (peak 1; Fig. 2, top left) and RpoN (peak 2; Fig. 2, top left). Two of the regulatory motifs in peak 1 are associated with iron metabolism and the motif in peak 2 is associated with nitrogen metabolism. Thus, this species’ local adaptation was likely in response to high bioavailable nitrogen and iron in the cecum compared to scarce levels of both in the rectum. Of note, iron-containing cofactors are required in many of the RpoN nitrogen metabolism pathways, suggesting that the 2 identified peaks represent a joint causal response to the divergent environment. Previous research has demonstrated that the various nutrient regimes throughout the gut have a significant impact on microbiome composition across a wide range of systems (21–23). Our results strongly suggest that spatial heterogeneity in nutrients, in addition to driving community composition, can also shape genetic variation within a microbial species in natural microbiota.

Beyond nutrient availability, the cecum and rectum differ markedly in environmental stability. The cecum represents a relatively stable niche, characterized by consistent nutrient input from the small intestine, prolonged transit time, strict anoxia, and a resident microbial community adapted to continuous fermentation (20). In contrast, the rectum is a more dynamic and fluctuating environment, defined by limited and variable nutrient availability, rapid transit, intermittent exposure to oxygen or the external environment, and periodic evacuation, all of which contribute to increased ecological instability (4,24,25). One possible mechanism by which microbial populations could adapt to such unstable and fluctuating environments is via mobile genetic elements (MGEs). MGEs can rapidly introduce nucleotide polymorphisms, chromosomal rearrangements, regulatory changes, and even entirely new genes, thereby increasing genetic variation within populations and facilitating swift adaptation to fluctuating environmental conditions (26–28). For example, previous research has highlighted the influence of MGEs in microbial adaptation to antibiotics (29) or in defense against phages (30). However, because the majority of MGE insertions are deleterious, for example by disrupting functional genes or regulatory pathways or promoting genome instability, microbial species have developed multiple mechanisms to prevent or mitigate the effects of MGEs (30,31).

Notably, we found significant local adaptation outliers in mobile genetic elements in three out of the ten species with outlier genes: *Cryptobacteroides* sp., *Faecousia* sp. A, *Phocaeicola sp017467385* (Fig. 2; Table S1); *Cryptobacteroides* sp., for example, displayed a single, extreme outlier peak located within a transposase gene (Fig.2, top right). Based on our results, we speculate that these species could be locally adapted for decreased MGE activity in the cecum, where the steady environment selects for genomic stability, and for increased MGE activity in the rectum, where the fluctuating environmental conditions select for increased genomic flexibility and adaptability. This would suggest that selection is strong enough to modulate MGE activity over very fine-grained spatial scales, balancing genomic stability and flexibility, and enabling microbial populations to thrive in the contrasting environments. Future experimental work is needed to validate the functional consequences of the identified MGE point mutations.

Having inferred the selection pressures likely driving local adaptation, we next considered how other forces might shape the adaptive process. Given that microbiome populations are suggested to experience powerful homogenizing forces, such as strong migration and clonal reproduction, how can local adaptation persist? One possibility is that the sampled microbiota populations were isolated between the cecum and rectum, which allowed genetic local adaptation to occur and persist without the counteracting force of migration. To this end, we examined potential signals of migration between populations by using intergenic diversity as a proxy for neutrally evolving regions. We found that cecum and rectum populations clustered more closely in ordination space within deer rather than between deer (Fig. S4), with significantly lower intergenic F_ST_ within than between deer (Fig. S5; p<0.001). These intergenic patterns indicate appreciable migration between microbial populations sampled from the same deer, suggesting that genetic local adaptation was able to occur in the face of migration.

We next examined the linked diversity surrounding our local adaptation outlier sites. Theory predicts that when selection is much stronger than migration, maladapted alleles will be purged by selection faster than migration can introduce them. This reduces the genetic diversity around locally adaptive sites (32–34). On the other hand, if migration is much stronger than selection (but not so strong as to collapse the locally adapted polymorphism), multiple allele classes can persist in a given environment at once, increasing the genetic diversity around locally adaptive sites (34,35). In our data, the linked diversity within populations at most local adaptation outliers showed elevated diversity relative to the genome-wide background (Fig. S6), suggesting that these species were locally adapting under strong incoming migration.

Given the evidence for substantial migration between populations, how are locally adapted populations able to maintain differentiation against a high influx of maladapted migrant genotypes and within the constraints of clonal reproduction? One possible explanation is that the species capable of genetic local adaptation were those with effective horizontal gene transfer (HGT). With effective HGT, a resident population could preserve locally adapted variants while at the same time allowing incoming migrants to rapidly acquire said variants. This dynamic could facilitate the development and maintenance of local adaptation, even against migration and clonal reproduction. To test this hypothesis, we computed the pairwise r² between all loci within a scaffold, normalized against the local genomic diversity, as a proxy for the homologous recombination rate. When we compared the recombination rate in 15 kb windows centered on local adaptation outliers to similar windows in the set of non-locally adapted species, we found that windows with local adaptation outliers exhibited significantly increased recombination rates (Fig. S7; p<0.001). The *Cryptobacteroides* sp. transposase local adaptation outlier, for example, is located in an extended ∼53 kb window of increased recombination (Fig. 3A), which was significantly enriched for mobile genetic elements in the assembly (Fig. 3B), including all necessary elements for an integrative and conjugative element. Furthermore, this region had a significantly different GC content (p<0.001) and codon usage (p<0.001) compared to the genomic background (Fig. 3C-D). Whereas the distinctive GC content and codon usage could potentially indicate misassembly, the absence of N’s within the 53 kb window and the surrounding 90 kb flanking nucleotides, coupled with a visual inspection of the aligned reads in IGV, revealed no evidence of assembly errors (Fig. S8). Instead, the elevated recombination rate and the high density and specific functions of the MGEs in this region strongly suggest its involvement in horizontal gene transfer, while the discordant GC content and codon usage patterns indicate a potential interspecific ancestral origin.

**Figure 3:**
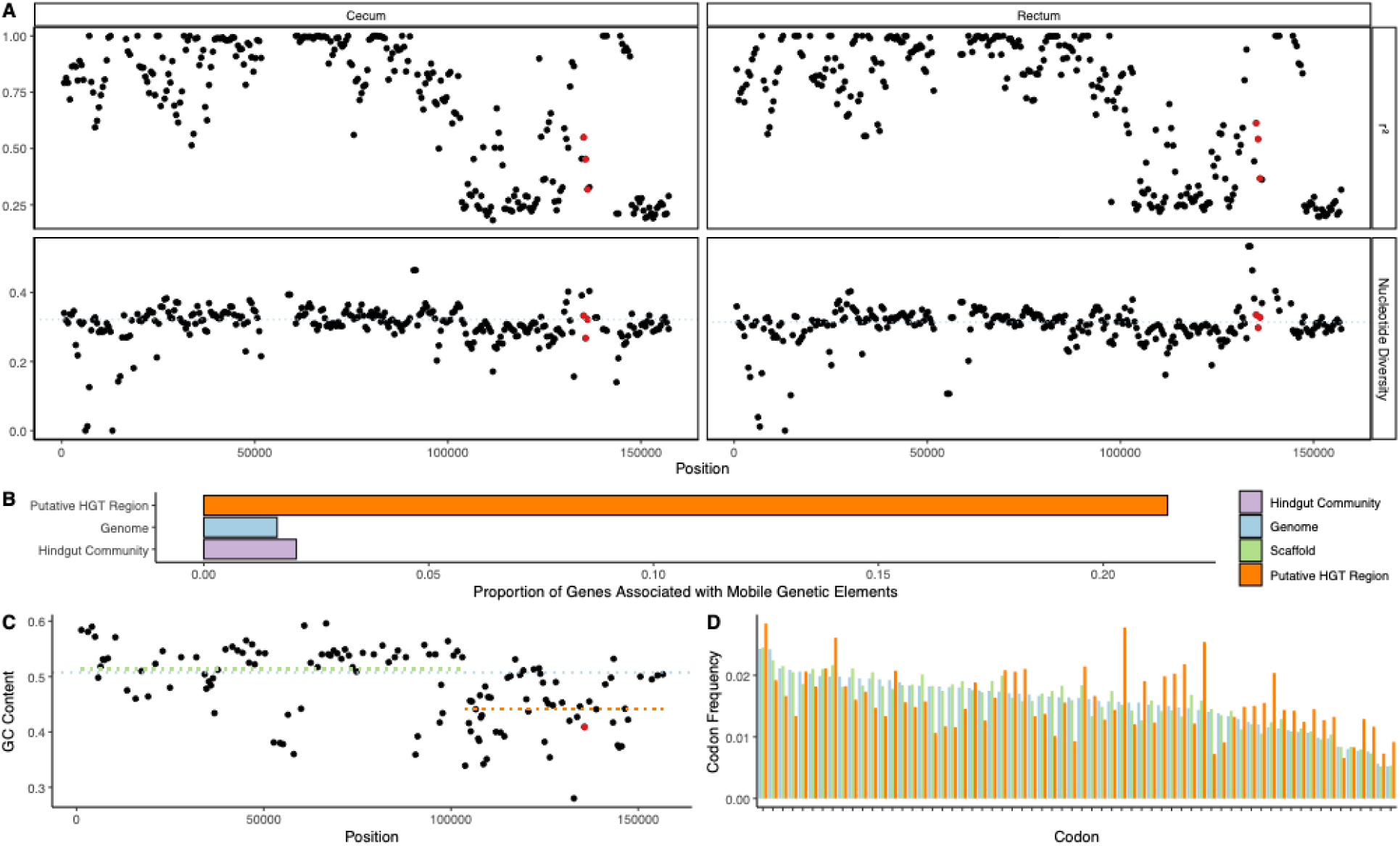
*Cryptobacteroides* sp. local adaptation outlier (transposase) lies in a 53k nucleotide window displaying significant signatures of horizontal gene transfer. (A) r^2^ as a proxy for homologous recombination rate, and nucleotide diversity in the cecum and rectum populations in 1,000 bp windows with 500 bp step size. (B) The proportion of genes associated with mobile genetic elements in the assemblies of the putative HGT window, the *Cryptobacteroides* sp. genome (ignoring the HGT window) and the entire hindgut community (ignoring *Cryptobacteroides* sp.). (C) GC content of all genes in the *Cryptobacteroides* sp. scaffold containing the putative HGT window. Dashed lines denote the mean GC for the HGT window (orange), the scaffold (ignoring the HGT window; green) and the genome (ignoring the scaffold; blue). (D) Frequency of each codon in the putative HGT window, the scaffold (ignoring the HGT window) and the genome (ignoring the scaffold). Windows (A) or the gene (C) corresponding to the putative local adaptation outlier transposase (Fig. 2, top right) are highlighted in red.

Our results offer compelling evidence that HGT plays an important role in driving genetic local adaptation in natural microbial populations, especially in high migration, high mixing regimes like the animal gastrointestinal tract. By allowing recent migrants to acquire clusters of coadapted alleles from the local population in a single event, HGT may enable more rapid adaptation than relying on *de novo* mutations, especially as the trait under selection is increasingly polygenic. The increased local recombination rates around local adaptation outliers, compared to similar regions in non-locally adapted species, suggest that a key difference between microbial species that are capable of genetic local adaptation and those that are not, lies in their respective recombination landscapes. In eukaryotes, both theory and empirical data support the idea that inheriting coadapted allele sets promotes local adaptation, often via chromosomal inversions that suppress recombination and preserve beneficial combinations across generations (36–41). This appears to contrast with our findings in microbes, where increased recombination via HGT is associated with local adaptation, however, it is possible that both strategies could reflect the same underlying principle: maintaining coadapted alleles in the face of migration. Thus, despite opposite patterns in recombination, mechanisms that facilitate local adaptation by promoting the inheritance of beneficial multi-locus genotypes could be universal across diverse life forms, shaped by the distinct genomic architectures and evolutionary constraints of each system.

Ultimately, our study reveals that genetic local adaptation is frequent in the hindgut microbiome, with 10/20 species displaying significant outliers between cecum and rectum populations across multiple deer. These findings challenge the common assumption that microbial community composition alone is sufficient to describe microbial function across different contexts. For example, the practice of generalizing results from fecal microbiome samples to different regions of the gut is widespread, likely because of accessibility and feasibility; however, we provide quantitative evidence calling for caution when interpreting results from such a method.

Our findings have direct implications for understanding microbial resilience and adaptability in response to other heterogeneous environments, for example, dietary changes, host immune pressures, or antimicrobial exposure. Just as microbial populations adapt to changes in nutrients or environmental stability, they may also undergo local adaptation to antimicrobials, allowing resistance alleles to be maintained in selective environments while they are being purged in others. The fine-grained spatial adaptation we observed suggests that antimicrobial resistance may similarly be shaped by the micro-environmental variation within compartments in the host, rather than solely by external antimicrobial exposure. Furthermore, we demonstrate that local adaptation and migration can potentially maintain locally adaptive polymorphisms within a single gut compartment, suggesting that a similar dynamic could maintain antimicrobial resistance alleles in compartments even when they are maladaptive, acting as a reservoir for future resistance. Overall, our findings underscore the fundamental role of genetic local adaptation in natural microbiome populations responding to environmental heterogeneity and highlight the need to consider the microbiome at both a genetic and spatial resolution.

## Materials and Methods

### Data Collection

We sampled gut microbiota from seven roe deer (*Capreolus capreolus*) from Brüggwald forest, Biel, Canton of Bern, Switzerland, on November 21, 2022. The sampling was performed directly after a regular hunting event, and all deer carcasses were sampled within <2 hours of shooting. We sampled each deer along its gastrointestinal tracts longitudinally at the same 10 predefined locations (Fig. 1A). For this, the entire gastrointestinal tract was extracted from the body. Samples were stored in sterile containers, transported under refrigerated conditions to the laboratory on the same day, and stored at -20°C until further processing.

### Extraction, Sequencing and Data Processing

We used the QIAamp PowerFecal Pro DNA Kit (Qiagen, Düsseldorf, Germany) to extract fecal DNA according to the manufacturer’s protocol. Briefly, we added a single fecal pellet to a PowerBead Pro Tube containing 800 µL of Solution CD1 and mixed it by vortexing. We homogenized the samples by bead beating at 30 Hz for 10 minutes using a tissue lyzer. We then centrifuged at 15,000 g for 1 minute and transferred about 500–600 µL of the supernatant into a clean 2 mL microcentrifuge tube. We added 200 µL of Solution CD2, vortexed, and centrifuged again at 15,000 g for 1 minute. We carefully transferred the resulting supernatant to a new tube, avoiding any pellets. Next, we added 600 µL of Solution CD3, vortexed, and loaded 650 µL of the lysate onto an MB Spin Column. We centrifuged at 15,000 g for 1 minute, discarded the flow-through, and applied the remaining lysate to the column. We washed the column sequentially with 500 µL each of Solution EA and Solution C5, centrifuging at 15,000 g for 1 minute after each wash. We performed a final centrifugation at 16,000 g for 2 minutes to remove the residual wash buffer. Finally, we eluted DNA with 50–100 µL of Solution C6 into a clean 1.5 mL elution tube by centrifugation at 15,000 g for 1 minute. We assessed DNA yield and purity using a NanoDrop spectrophotometer and stored the samples at −20°C until further use. DNA samples were then sequenced on an Illumina Novaseq (150 bp PE mode) at the next-generation sequencing facility of the University of Bern, Switzerland. We trimmed adaptors and low-quality bases with fastp v0.19.5 (42) and removed PCR duplicates with FastUniq v1.1 (43).

### *De novo* Assembly, Binning, and Dereplication

We performed *de novo* assembly using metaSPAdes v3.15.3 (44) on metagenomic mode (--meta) and with four different k-mer lengths (21, 33, 55, 77 bp). We filtered the resulting assemblies based on scaffold size (≥1,500 bp) and k-mer coverage (≥2x) of the largest (77 bp) k-mer. We used Bowtie2 v2.4.4 (45) to align the processed reads against our filtered assemblies and SAMtools (46) to sort and index them.

We used three different binning tools to bin sequences into putative single-species genomes, MetaBAT2 v2.16 (47), MaxBin2 v2.2.7 (48), and CONCOCT v1.1.0 (49). Additionally, we used Binning_refiner (50) to generate hybrid bin sets in all pairwise combinations and the triplewise combination (e.g., merge highly similar bins, remove likely contamination, reassign putatively misclassified contigs). We then used CheckM2 v1.0.2 (51) to assess the quality of each bin (e.g. completeness, contamination, GC content

We filtered all bins by completeness (≥70%) and contamination (≤10%) and then used custom scripts from metaWRAP (52) to compare analogous bins across all binning strategies and choose the best bin (consolidate_two_sets_of_bins.py), and to dereplicate any contigs that were duplicated over the remaining bins (dereplicate_contigs_in_bins.py). Finally, we took the entire set of genomes (i.e., all deer, all compartments) and dereplicated them using dRep v3.5.0 (53), based on primary and secondary clusters of 80% and 95% ANI respectively. We chose the best genome out of all replicate genomes based on completeness, contamination, strain heterogeneity, N50, and centrality.

### Taxonomy Classification and Quantification of Relative Abundance

We inferred the taxonomical classification of our final genomes using GTDB-tk v2.4.0 release 220 (54). We used Salmon v1.10.2 (55) on metagenomic mode (--meta) to estimate the scaffold abundances in each sample. To estimate the abundance of a given species in a sample, we summed the total number of nucleotides in all scaffolds corresponding to the species and divided by the size of its assembled genome. Nucleotide abundances were transformed into relative abundances by scaling them by the total number of nucleotides in the entire sample. In order to perform principal component analyses on species relative abundances, we first applied centered log-ratio transformations.

### Variant Calling

To call SNPs, we used inStrain v1.9.0 (56) with default settings, i.e., a variant was called if there was at least 5 depth at the position, at least 5% variant frequency, and the variant depth was greater than a null model based on the overall depth at the locus. To capture sites that were fixed for the reference allele in one compartment and for a variant in another (and thus not called by the above procedure), we merged BAM files from different compartments using SAMtools, called SNPs on the merged BAMs (≥10 depth, ≥2.5% variant frequency) and then back-calculated the base calls at each locus in the original compartment. Finally, we filtered out any new SNPs called from the merged BAM files that were not at least 5% variant frequency in one compartment (e.g., a SNP at 2.5% variant frequency in both compartments would be filtered out).

### Outlier Identification

To identify outlier SNPs, we first filtered our called SNPs down to include only SNPs called in both compartments in at least two deer (replicates) for a given species. We then focused our analysis on the top 20 species by filtered SNP counts for our downstream outlier analyses. To identify SNPs putatively responding to selection, we performed Cochran-Mantel-Haenszel tests on allele counts at each SNP to find sites with the largest, concordant allele frequency changes between compartments across multiple deer. For all SNPs within a given species, we used the Benjamini-Hochberg procedure to correct for false discovery rates at a threshold of 0.05. We then classified all significant SNPs that were also in the top 0.01% most extreme SNPs for a given microbial species as outliers.

We classified any gene with an outlier SNP in it as an outlier gene. We identified intergenic outliers as intergenic outlier SNPs that were in or within 50 bp of motifs that significantly matched experimentally validated regulatory motifs. We used FIMO (57) and the CollecTF database (58) to identify regulatory motifs. Notably, with the described method any gene or intergenic window with (potentially) a single outlier SNP in it was classified as an outlier, in essence ignoring the linkage relationships of the nearby region. We structured our outlier detection in this manner to improve our sensitivity to detecting outliers from *de novo* mutation and soft sweeps.

### Outlier Functions

We used Prodigal v2.6.3 (59) to infer the gene space in each genome and eggNOG v2.0.1 (60) or GAIA (61) to infer gene functions. For intergenic outliers, we assumed similar functions for regulatory regions as reported in MycoBrowser (17), KEGG (18), or EcoCyc (19) databases, or in the primary literature.

### Additional Analyses

We calculated F_ST_ according to (62), and nucleotide diversity according to (63). We used InStrain to calculate r^2^ and normalized r^2^ between loci within a scaffold as proxies for homologous recombination rate for comparison within and between species, respectively. We identified genes associated with mobile genetic elements as those with the following keywords in their inferred functions: capsid, competence, conjugation, conjugative, DNA uptake, insertion sequence, integrase, IS element, mobilizable, phage, plasmid, prophage, recombinase, relaxase, resolvase, tail fiber, terminase, transposase, transposon, type III secretion/secretory, type IV pilus, type IV secretion/secretory, type VI secretion/secretory, viral, Mob[A:Z], Par[A:Z], Rep[A:Z], Tra[A:Z].

### Statistical Testing

To test for matches between our outlier regulatory regions (n=9) and regulatory motifs of known function we used likelihood ratio tests. In order to test for a difference in intergenic F_ST_ between cecum and rectum populations *within* a deer versus *between* deer, we used a permutation test with species as a block factor. Because the original distributions were quite different, we used a Kolmogorov–Smirnov permutation test to test for a difference in r^2^ (normalized) between outlier windows and background windows. We used a permutation test to test for a difference in GC content between the putative HGT window in *Cryptobacteroides* sp. and the genome background. To test for different codon usage or mobile genetic element enrichment in our putative HGT window compared to the background, we used a chi-squared test and Fisher’s exact test, respectively. We used a PERMANOVA with the gut region (foregut, midgut, hindgut) and the individual deer as explanatory factors to test for differences between microbial communities.

As previously mentioned, we corrected for the number of SNPs in a given species during our outlier detection by applying a Benjamini-Hochberg procedure (FDR=0.05). Beyond our SNP outlier detection, we also performed 15 other significance tests (9 regulatory motifs, 6 others). Therefore, we performed a study-wide Bonferroni correction resulting in a study-wide ɑ value on the order of 0.001.

### Methods External Dependencies

The following were external dependencies for the primary tools used in our study.

| Tool |
| --- |
| DIAMOND (64) |
| FastANI (65) |
| FastTree 2 (66) |
| FragGeneScan (67) |
| HMMER (68) |
| Mash (69) |
| MMseqs2 (70) |
| pplacer (71) |
| skani (72) |

## Author contributions

Conceptualization: CB, JB, BY

Methodology: RJJ, AK, SP, BY, JB, CB

Data Analysis: RJJ

Visualization: RJJ

Funding acquisition: CB, JB, BY, RJJ

Supervision: CB, SP, AK

Writing – original draft: RJJ

Writing – review & editing: RJJ, AK, SP, BY, JB, CB

## Data Availability

Code is available at github.com/banklab/Russ_Metagenomics and will be archived on Zenodo upon publication of the paper. Data is deposited at the European Bioinformatics Institute with an embargo period.

## Acknowledgements

We thank Daniel Trachsel for generously providing access to the hunting grounds on the day of the hunt and for supplying the deer carcasses essential to this study. We thank Belinda Köchle for support with the surgical procedures involved in sample collection and Nicole Nesvadba for expertise in DNA extraction and wet lab. We thank Katie Peichel and the Theoretical Ecology and Evolution lab for helpful comments and discussion. We thank the UBELIX () and IBU HPC clusters at the University of Bern for providing computational resources and the Next Generation Sequencing Platform at the University of Bern for performing the high-throughput sequencing.

This project was supported by NSERC PGSD3-546782-2020 to R.J., SNSF Grant 10001034 to S.P., SNSF Starting Grant TMSGI3_211300 to B.Y., Novartis Foundation for medical-biological Research Grant 22A018 to J.B., and SNSF Grant 315230_204838/1 to C.B.. Figure 1 image (AdobeStock_530047064) was used with permission under Adobe’s Education License.

## Supplementary Figures and Tables

**Figure S1:**
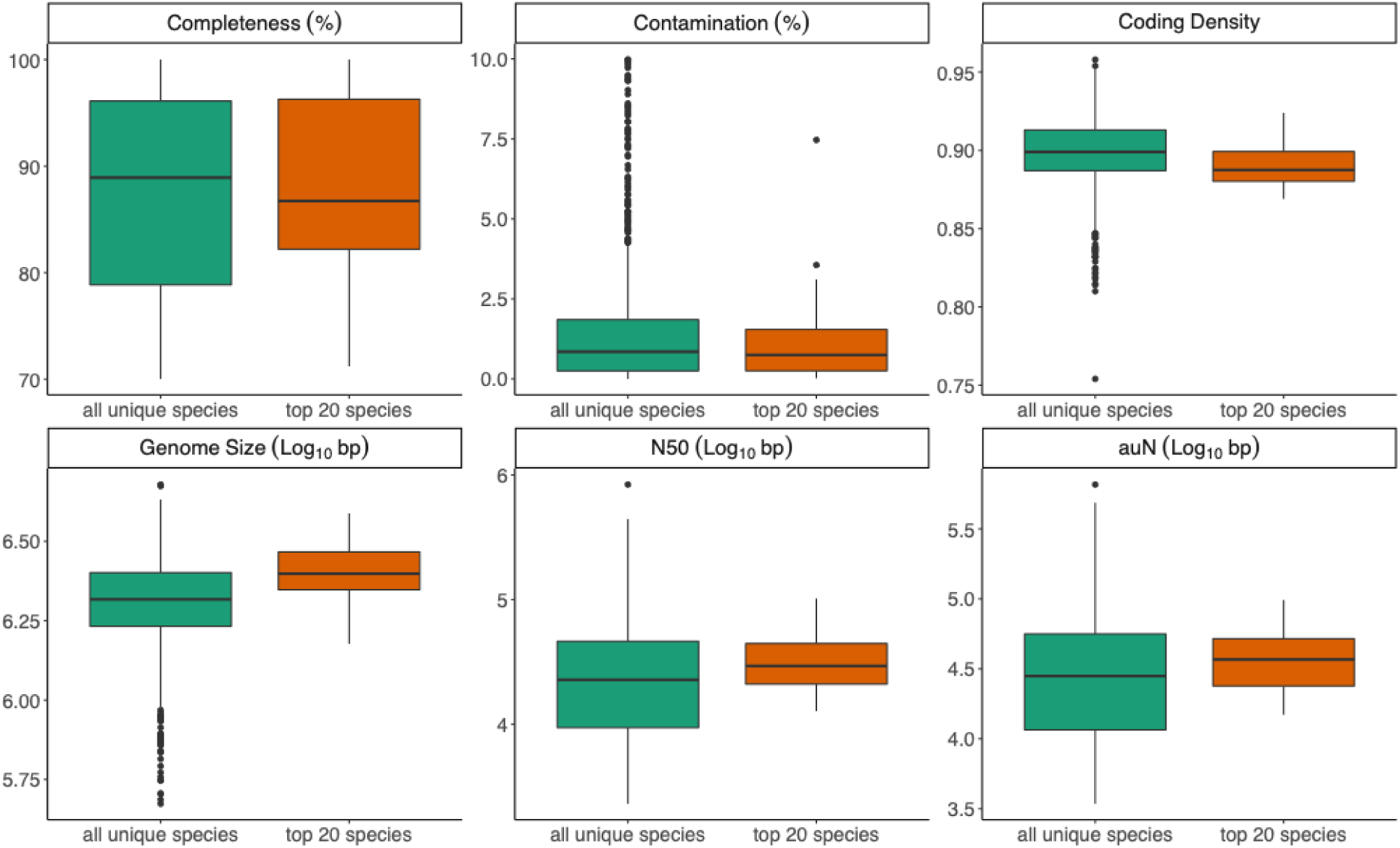
Assembly metrics for all unique species (909 species) and the top 20 species (based on SNP volume). The facets denote completeness, the estimated percentage of the genome that is present in the assembly; contamination, the estimated percentage of foreign DNA in the assembly; coding density, the percentage of protein coding sequences in the assembly; genome size, the size of the assembly; N50, the length of the shortest scaffold at 50% of the total assembly size (i.e., contiguity); auN, area under the Nx curve (i.e., contiguity).

**Figure S2:**
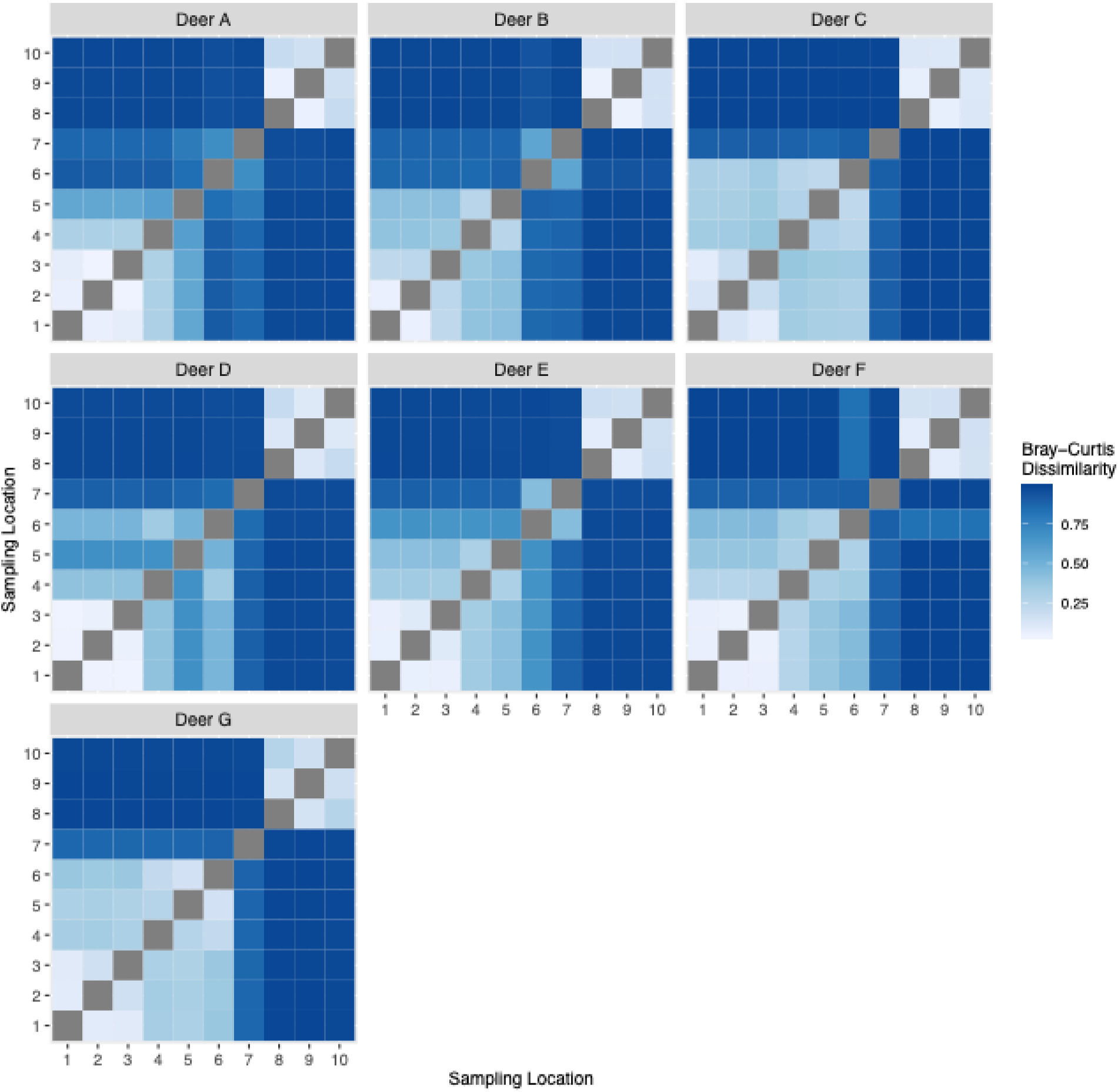
Bray-Curtis dissimilarity indices for species abundances between all sampling locations within a given deer indicate 2-3 main community types along the intestinal tract, where compartments 1-5, and 8-10 cluster across all seven deer. Sampling locations are described as in Fig. 1.

**Figure S3:**
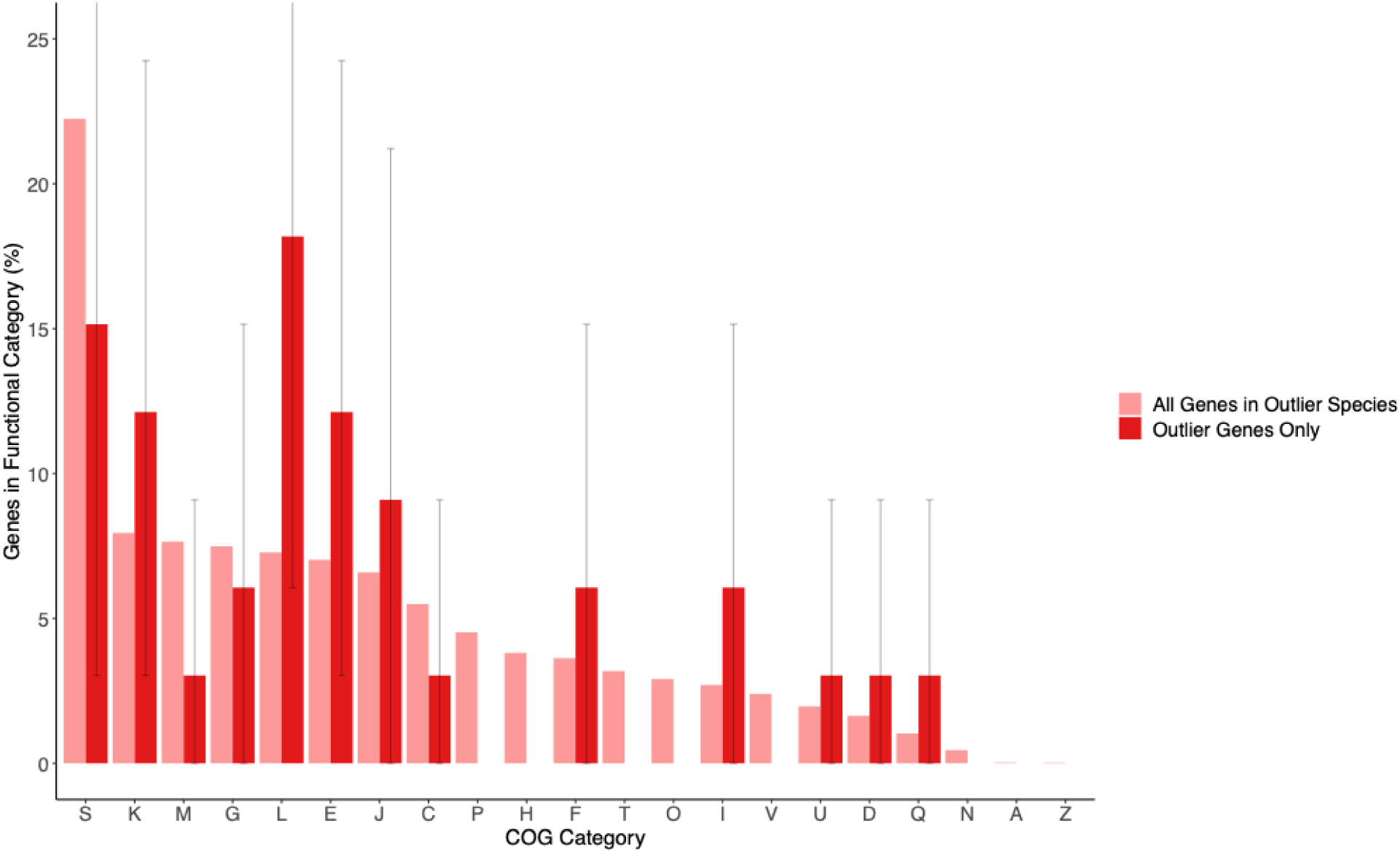
Distribution of gene functions in the 10 putatively locally adapted species for local adaptation outlier genes and for all genes. Error bars denote 95% confidence intervals. Gene functions are ascribed according to the COG database (13). Note that functional categories are not mutually exclusive for a given gene. COG categories: S, Function Unknown; K, Transcription; M, Cell Wall/Membrane/Envelope Biogenesis; G, Carbohydrate Metabolism and Transport; L, Replication, Recombination, and Repair; E, Amino Acid Metabolism and Transport; J, Translation, Ribosomal Structure, and Biogenesis; C, Energy Production and Conversion; P, Inorganic Ion Transport and Metabolism; H, Coenzyme Metabolism; F, Nucleotide Metabolism and Transport; T, Signal Transduction Mechanisms; O, Posttranslational Modification, Protein Turnover, Chaperones; I, Lipid Metabolism; V, Defense Mechanisms; U, Intracellular Trafficking, Secretion, and Vesicular Transport; D, Cell Cycle Control, Cell Division, Chromosome Partitioning; Q, Secondary metabolites biosynthesis, transport, and catabolism; N, Cell Motility; A, RNA Processing and Modification; Z, Cytoskeleton.

**Figure S4:**
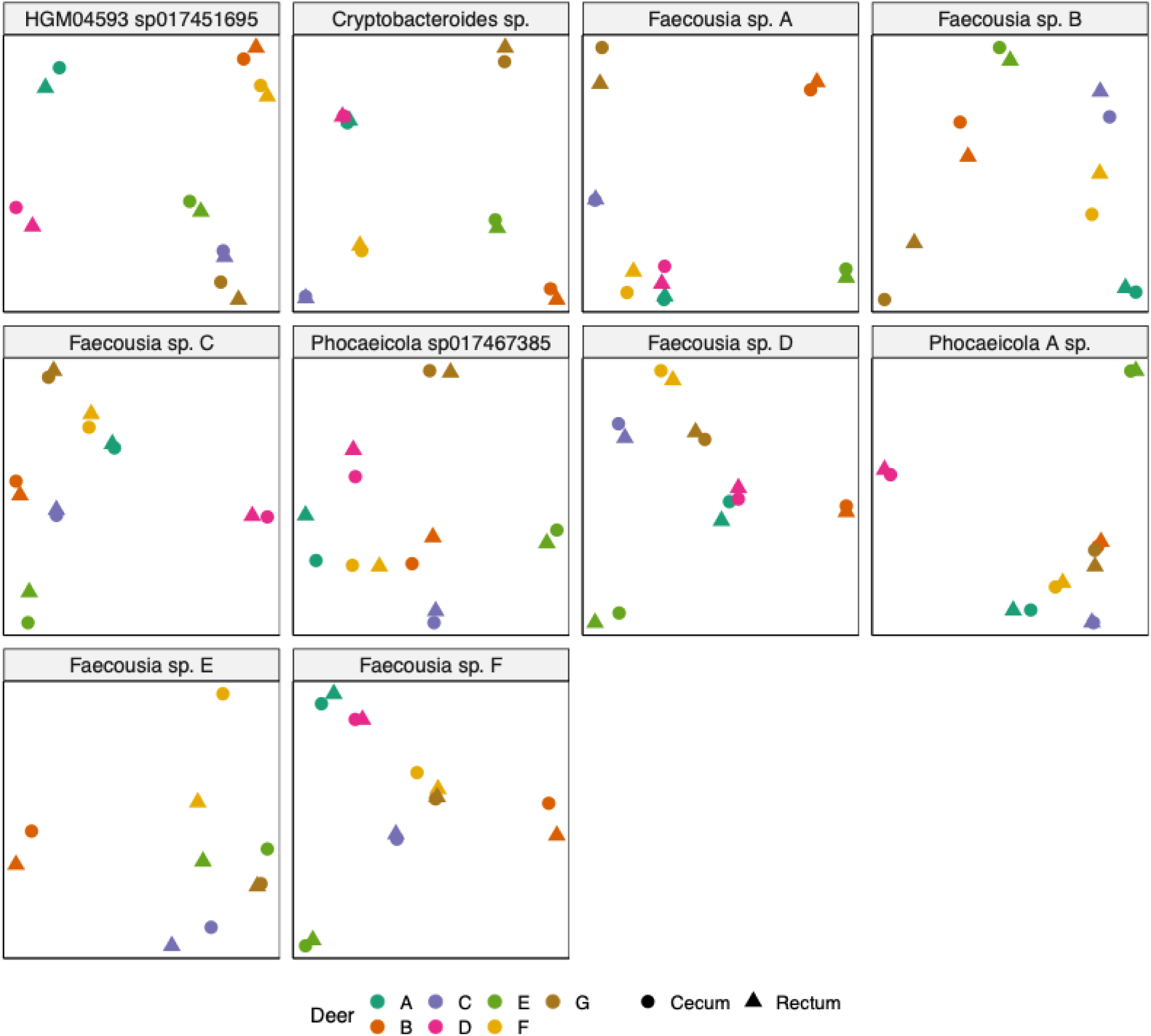
Non-metric multidimensional scaling on intergenic allele frequencies of the 10 putatively locally adapted species suggest migration between populations within deer. In general, populations within the same deer clustered more closely together in ordination space than populations between deer, suggesting migration within deer.

**Figure S5:**
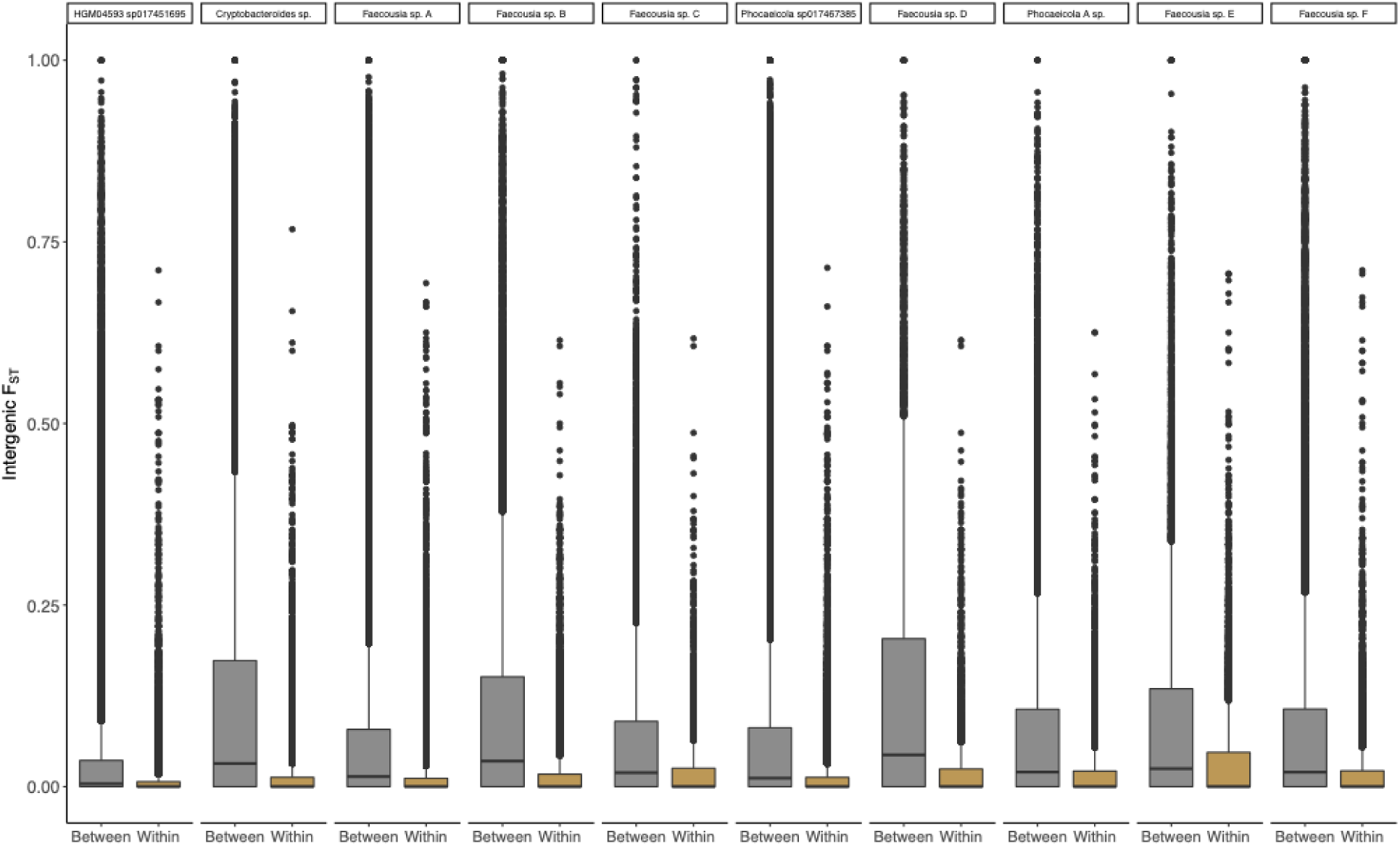
Intergenic F_ST_ between cecum and rectum populations between and within deer for the 10 putatively locally adapted species suggests migration between populations within deer. Each point represents a SNP.

**Figure S6:**
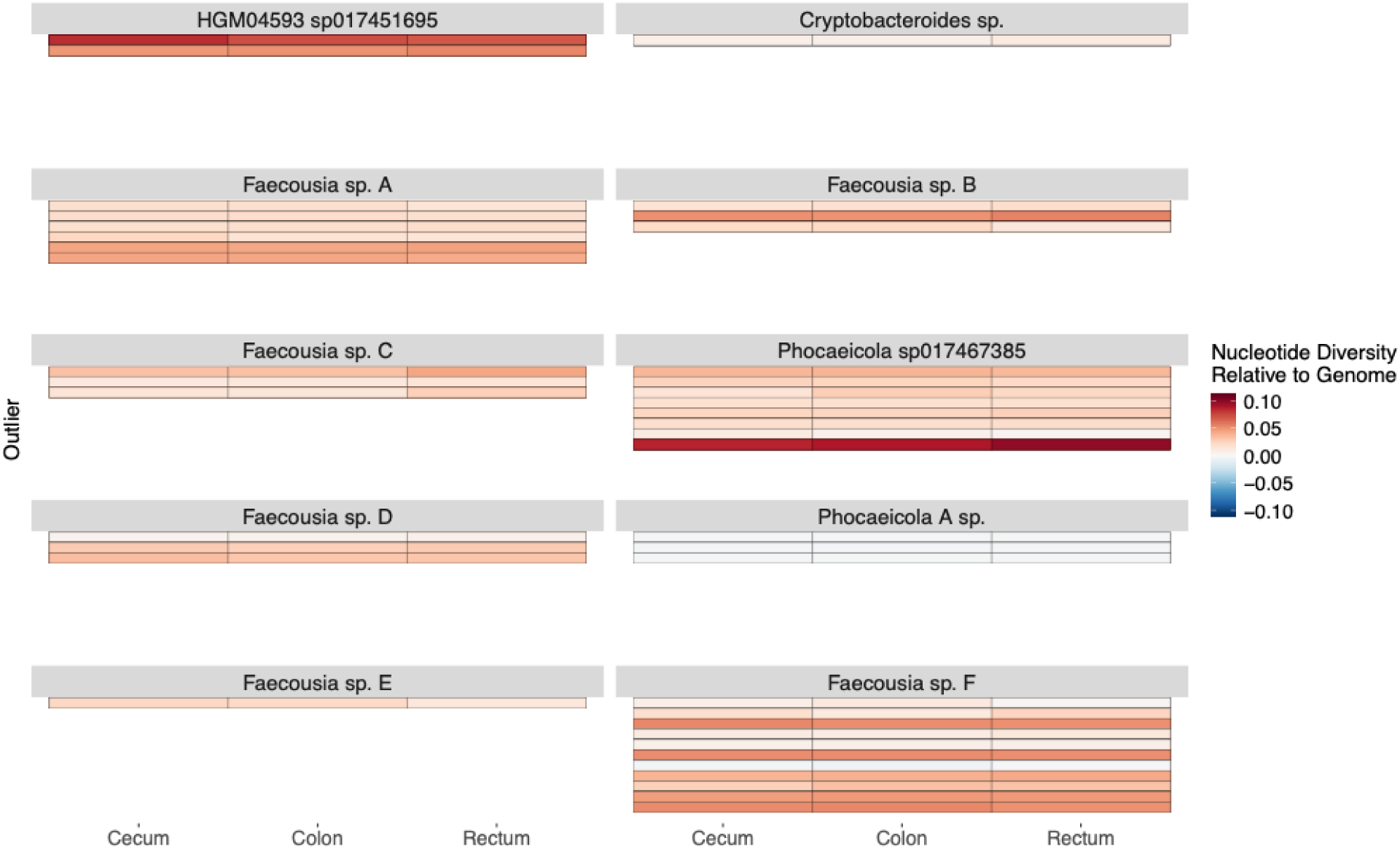
Nucleotide diversity tends to be elevated at outlier genes or outlier intergenic windows compared to the genome background. Diversity is shown as the difference between an outlier and the genome-wide background for the cecum, colon, and rectum populations, sampling locations 8-10 respectively (Fig. 1).

**Figure S7:**
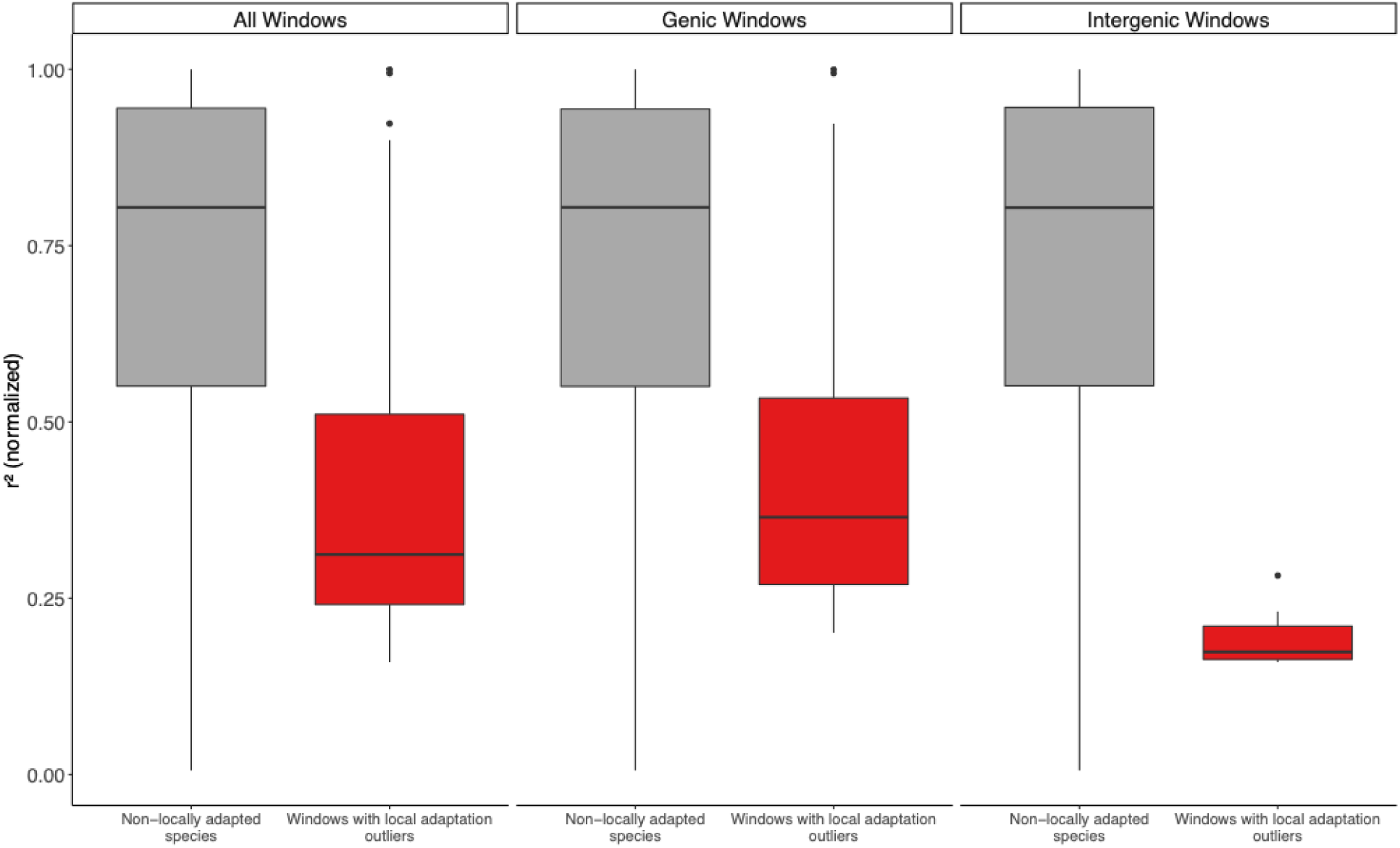
Normalized r^2^ was reduced in windows containing local adaptation outlier genes compared to analogous windows in non-locally adapted species. r^2^ (normalized by the local genomic diversity) in 15 kbp windows for the top 20 species (by SNP volume). All windows in the 10 putatively non-locally adapted species (grey) or windows centered on local adaptation outliers in the 10 putatively locally adapted species (red).

**Figure S8:**
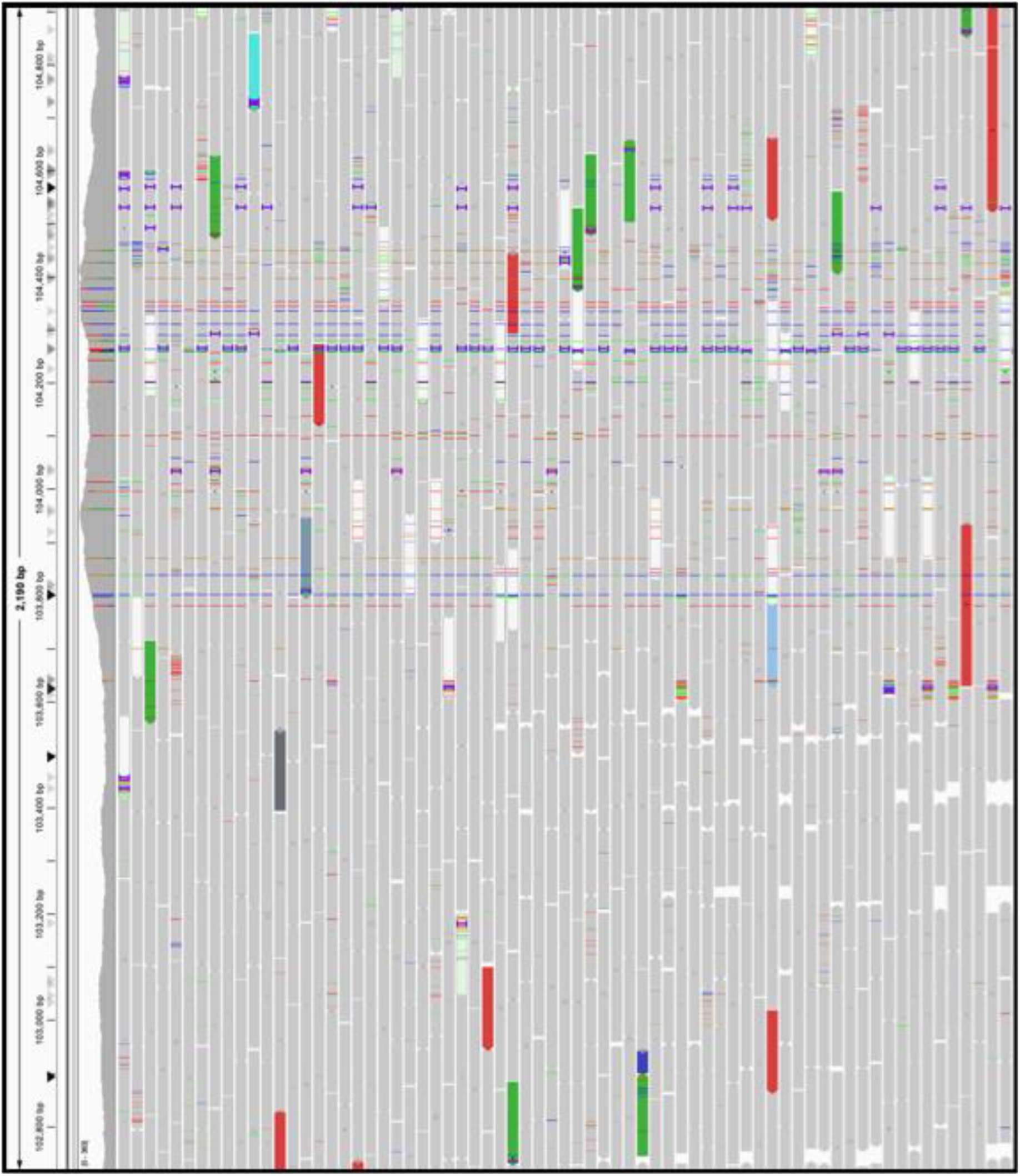
Snapshot of IGV from *Cryptobacteroides sp.* alignment. We inspected this region for evidence of misassembly. Note that the snapshot is truncated in both dimensions (i.e., genome length and alignment depth).

**Table S1:** Genic and intergenic outliers for the 10 species showing putative genetic local adaptation between cecum and rectum. The predicted functions of the genic outliers are described according to the function of the closest known orthologs; the predicted function of the intergenic outliers are described according to the function of significantly matching (LRT, α=0.001) regulatory motifs within 50 bp of the outlier SNP.

| Species | Scaffold | Genomic Element | Description (gene)<br>Motif (intergenic) | Biological Function |
| --- | --- | --- | --- | --- |
| *HGM04593 sp017451695* | 24575 | Intergenic | Fur (5 bp away) | iron metabolism |
| *HGM04593 sp017451695* | 24575 | Intergenic | PvdS | pyoverdine synthesis |
| *HGM04593 sp017451695* | 24575 | Intergenic | DosR | hypoxia, dormancy |
| *HGM04593 sp017451695* | 40195 | Intergenic | RpoN | nitrogen metabolism |
| *Cryptobacteroides* sp. | 16 | Gene | Transposase | replication, recombination and repair |
| *Faecousia* sp. A | 2096 | Gene | Phage integrase, N-terminal SAM-like domain | replication, recombination and repair |
| *Faecousia* sp. A | 3589 | Gene | NA | NA |
| *Faecousia* sp. A | 3589 | Gene | Psort location<br>CytoplasmicMembrane, score | no functional prediction |
| *Faecousia* sp. A | 21682 | Intergenic | MatP (9 bp away) | chromosomal segregation, cell division |
| *Faecousia* sp. A | 51206 | Intergenic | CtrA (46 bp away) | cell division |
| *Faecousia* sp. A | 51206 | Intergenic | HgtR (25 bp away) | energy metabolism |
| *Faecousia* sp. B | 216 | Gene | Type IV secretory pathway, VirB4 components | intracellular trafficking, secretion, and vesicular transport |
| *Faecousia* sp. B | 2061 | Gene | Catalyzes the phosphorylation of D-fructose 6-phosphate, the first committing step of glycolysis. Uses inorganic phosphate (PPi) as phosphoryl donor instead of ATP like common ATP-dependent phosphofructokinases (ATP-PFKs), which renders the reaction reversible, and can thus function both in glycolysis and gluconeogenesis. Consistently, PPi-PFK can replace the enzymes of both the forward (ATP-PFK) and reverse (fructose-bisphosphatase (FBPase)) reactions | carbohydrate metabolism and transport |
| *Faecousia* sp. B | 4770 | Gene | BadF/BadG/BcrA/BcrD ATPase family | lipid metabolism |
| *Faecousia* sp. C | 1334 | Gene | DNA-templated transcription, initiation | transcription |
| *Faecousia* sp. C | 2526 | Gene | NifU-like N terminal domain | energy production and conversion |
| *Faecousia* sp. C | 1334 | Intergenic | CRP | energy metabolism |
| *Phocaeicola* sp017467385 | 123 | Gene | Transposase, mutator | replication, recombination and repair |
| *Phocaeicola* sp017467385 | 1023 | Gene | transposase, IS4 | replication, recombination and repair |
| *Phocaeicola* sp017467385 | 34868 | Gene | COG2003 DNA repair | replication, recombination and repair |
| *Phocaeicola* sp017467385 | 37392 | Gene | Belongs to the 'phage' integrase family | replication, recombination and repair |
| *Phocaeicola* sp017467385 | 43588 | Gene | Helix-turn-helix domain | transcription |
| *Phocaeicola* sp017467385 | 48904 | Gene | Psort location Cytoplasmic, score 8.96 | no functional prediction |
| *Phocaeicola sp017467385* | 58528 | Gene | D-alanine [D-alanyl carrier protein] ligase activity | secondary metabolites biosynthesis, transport and catabolism |
| *Phocaeicola sp017467385* | 60380 | Intergenic | HrpX (39 bp away) | NA |
| *Faecousia* sp. D | 1042 | Gene | Catalyzes the reversible phosphatidyl group transfer from one phosphatidylglycerol molecule to another to form cardiolipin (CL) (diphosphatidylglycerol) and glycerol | lipid metabolism |
| *Faecousia* sp. D | 4636 | Gene | Endoribonuclease that initiates mRNA decay | cell wall structure and biogenesis and outer membrane |
| *Faecousia* sp. D | 6454 | Gene | Permease family | no functional prediction |
| *Phocaeicola A* sp. | 38 | Gene | Psort location<br>CytoplasmicMembrane, score 10.00 | no functional prediction |
| *Phocaeicola A* sp. | 822 | Gene | GXGXG motif | amino acid metabolism and transport |
| *Phocaeicola A* sp. | 3424 | Gene | Lytic transglycosylase with a strong preference for naked glycan strands that lack stem peptides | cell division and chromosome partitioning |
| *Faecousia* sp. E | 5489 | Gene | Catalyzes the ATP-dependent phosphorylation of L-homoserine to L-homoserine phosphate | amino acid metabolism and transport |
| *Faecousia* sp. F | 144 | Gene | Psort location<br>CytoplasmicMembrane, score | no functional prediction |
| *Faecousia* sp. F | 209 | Gene | Transcriptional regulator | transcription |
| *Faecousia* sp. F | 308 | Gene | Involved in mRNA degradation. Catalyzes the phosphorolysis of single-stranded polyribonucleotides processively in the 3'- to 5'- direction | translation, including ribosome structure and biogenesis |
| *Faecousia* sp. F | 369 | Gene | Threonine synthase N terminus | amino acid metabolism and transport |
| *Faecousia* sp. F | 935 | Gene | CobB/CobQ-like glutamine amidotransferase domain | nucleotide metabolism and transport |
| *Faecousia* sp. F | 1203 | Gene | PEP-utilising enzyme, TIM barrel domain | carbohydrate metabolism and transport |
| *Faecousia* sp. F | 2877 | Gene | tRNA (Uracil-5-)-methyltransferase | translation, including ribosome structure and biogenesis |
| *Faecousia* sp. F | 3559 | Gene | Belongs to the formate-tetrahydrofolate ligase family | nucleotide metabolism and transport |
| *Faecousia* sp. F | 3559 | Gene | Belongs to the class-II aminoacyl-tRNA synthetase family | translation, including ribosome structure and biogenesis |
| *Faecousia* sp. F | 3696 | Gene | DNA-dependent RNA polymerase catalyzes the transcription of DNA into RNA using the four ribonucleoside triphosphates as substrates | transcription |
| *Faecousia* sp. F | 7716 | Gene | Glutamate/Leucine/Phenylalanine/Valine dehydrogenase | amino acid metabolism and transport |

## References

1. Hou K, Wu ZX, Chen XY, Wang JQ, Zhang D, Xiao C, et al. Microbiota in health and diseases. Signal Transduct Target Ther. 2022 Apr 23;7(1):135.

2. Crespo-Piazuelo D, Estellé J, Revilla M, Criado-Mesas L, Ramayo-Caldas Y, Óvilo C, et al. Characterization of bacterial microbiota compositions along the intestinal tract in pigs and their interactions and functions. Sci Rep. 2018 Aug 24;8(1):12727.

3. Wan XL, McLaughlin RW, Zheng JS, Hao YJ, Fan F, Tian RM, et al. Microbial communities in different regions of the gastrointestinal tract in East Asian finless porpoises (Neophocaena asiaeorientalis sunameri). Sci Rep. 2018 Sep 20;8(1):14142.

4. Anders JL, Moustafa MAM, Mohamed WMA, Hayakawa T, Nakao R, Koizumi I. Comparing the gut microbiome along the gastrointestinal tract of three sympatric species of wild rodents. Sci Rep. 2021 Oct 7;11(1):19929.

5. Tong F, Wang T, Gao NL, Liu Z, Cui K, Duan Y, et al. The microbiome of the buffalo digestive tract. Nat Commun. 2022 Feb 10;13(1):823.

6. Dubinsky V, Dotan I, Gophna U. Strains Colonizing Different Intestinal Sites within an Individual Are Derived from a Single Founder Population. Sansonetti PJ, editor. mBio. 2023 Feb 28;14(1):e03456–22.

7. Wasney M, Briscoe L, Wolff R, Ghezzi H, Tropini C, Garud N. Uniform bacterial genetic diversity along the guts of mice inoculated with human stool [Internet]. 2025 [cited 2025 Jun 4]. Available from: http://biorxiv.org/lookup/doi/10.1101/2025.01.28.635365

8. Yilmaz B, Mooser C, Keller I, Li H, Zimmermann J, Bosshard L, et al. Long-term evolution and short-term adaptation of microbiota strains and sub-strains in mice. Cell Host Microbe. 2021 Apr;29(4):650–663.e9.

9. Garud NR, Good BH, Hallatschek O, Pollard KS. Evolutionary dynamics of bacteria in the gut microbiome within and across hosts. Gordo I, editor. PLOS Biol. 2019 Jan 23;17(1):e3000102.

10. Zhao S, Lieberman TD, Poyet M, Kauffman KM, Gibbons SM, Groussin M, et al. Adaptive Evolution within Gut Microbiomes of Healthy People. Cell Host Microbe. 2019 May;25(5):656–667.e8.

11. Zahavi L, Lavon A, Reicher L, Shoer S, Godneva A, Leviatan S, et al. Bacterial SNPs in the human gut microbiome associate with host BMI. Nat Med. 2023 Nov;29(11):2785–92.

12. Valles-Colomer M, Blanco-Míguez A, Manghi P, Asnicar F, Dubois L, Golzato D, et al. The person-to-person transmission landscape of the gut and oral microbiomes. Nature. 2023 Feb 2;614(7946):125–35.

13. Tatusov RL. The COG database: a tool for genome-scale analysis of protein functions and evolution. Nucleic Acids Res. 2000 Jan 1;28(1):33–6.

14. Leoni L, Orsi N, De Lorenzo V, Visca P. Functional Analysis of PvdS, an Iron Starvation Sigma Factor of *Pseudomonas aeruginosa*. J Bacteriol. 2000 Mar 15;182(6):1481–91.

15. Laub MT, Chen SL, Shapiro L, McAdams HH. Genes directly controlled by CtrA, a master regulator of the *Caulobacter* cell cycle. Proc Natl Acad Sci. 2002 Apr 2;99(7):4632–7.

16. Ueki T, Lovley DR. Genome-wide gene regulation of biosynthesis and energy generation by a novel transcriptional repressor in Geobacter species. Nucleic Acids Res. 2010 Jan;38(3):810–21.

17. Kapopoulou A, Lew JM, Cole ST. The MycoBrowser portal: A comprehensive and manually annotated resource for mycobacterial genomes. Tuberculosis. 2011 Jan;91(1):8–13.

18. Kanehisa M, Sato Y, Kawashima M, Furumichi M, Tanabe M. KEGG as a reference resource for gene and protein annotation. Nucleic Acids Res. 2016 Jan 4;44(D1):D457–62.

19. Moore LR, Caspi R, Boyd D, Berkmen M, Mackie A, Paley S, et al. Revisiting the y-ome of *Escherichia coli*. Nucleic Acids Res. 2024 Nov 11;52(20):12201–7.

20. Gressley TF, Hall MB, Armentano LE. RUMINANT NUTRITION SYMPOSIUM: Productivity, digestion, and health responses to hindgut acidosis in ruminants1. J Anim Sci. 2011 Apr 1;89(4):1120–30.

21. Donaldson GP, Lee SM, Mazmanian SK. Gut biogeography of the bacterial microbiota. Nat Rev Microbiol. 2016 Jan;14(1):20–32.

22. Pereira FC, Berry D. Microbial nutrient niches in the gut. Environ Microbiol. 2017 Apr;19(4):1366–78.

23. Tropini C, Earle KA, Huang KC, Sonnenburg JL. The Gut Microbiome: Connecting Spatial Organization to Function. Cell Host Microbe. 2017 Apr;21(4):433–42.

24. Jensen BAH, Heyndrickx M, Jonkers D, Mackie A, Millet S, Naghibi M, et al. Small intestine vs. colon ecology and physiology: Why it matters in probiotic administration. Cell Rep Med. 2023 Sep;4(9):101190.

25. Yilmaz B, Macpherson AJ. Delving the depths of ‘terra incognita’ in the human intestine — the small intestinal microbiota. Nat Rev Gastroenterol Hepatol. 2025 Jan;22(1):71–81.

26. Casacuberta E, González J. The impact of transposable elements in environmental adaptation. Mol Ecol. 2013 Mar;22(6):1503–17.

27. Schrader L, Schmitz J. The impact of transposable elements in adaptive evolution. Mol Ecol. 2019 Mar;28(6):1537–49.

28. Pimpinelli S, Piacentini L. Environmental change and the evolution of genomes: Transposable elements as translators of phenotypic plasticity into genotypic variability. Herrel A, editor. Funct Ecol. 2020 Feb;34(2):428–41.

29. Partridge SR, Kwong SM, Firth N, Jensen SO. Mobile Genetic Elements Associated with Antimicrobial Resistance. Clin Microbiol Rev. 2018 Oct;31(4):e00088–17.

30. Chevallereau A, Westra ER. Bacterial immunity: Mobile genetic elements are hotspots for defence systems. Curr Biol. 2022 Sep;32(17):R923–6.

31. Schmitz M, Querques I. DNA on the move: mechanisms, functions and applications of transposable elements. FEBS Open Bio. 2024 Jan;14(1):13–22.

32. Nordborg M. Structured Coalescent Processes on Different Time Scales. Genetics. 1997 Aug 1;146(4):1501–14.

33. Aeschbacher S, Bürger R. The Effect of Linkage on Establishment and Survival of Locally Beneficial Mutations. Genetics. 2014 May 1;197(1):317–36.

34. Jasper RJ, Yeaman S. Local adaptation can cause both peaks and troughs in nucleotide diversity within populations.

35. Charlesworth B, Nordborg M, Charlesworth D. The effects of local selection, balanced polymorphism and background selection on equilibrium patterns of genetic diversity in subdivided populations. Genet Res. 1997 Oct;70(2):155–74.

36. Kirkpatrick M, Barton N. Chromosome Inversions, Local Adaptation and Speciation. Genetics. 2006 May 1;173(1):419–34.

37. Lowry DB, Willis JH. A Widespread Chromosomal Inversion Polymorphism Contributes to a Major Life-History Transition, Local Adaptation, and Reproductive Isolation. Barton NH, editor. PLoS Biol. 2010 Sep 28;8(9):e1000500.

38. Joron M, Frezal L, Jones RT, Chamberlain NL, Lee SF, Haag CR, et al. Chromosomal rearrangements maintain a polymorphic supergene controlling butterfly mimicry. Nature. 2011 Sep;477(7363):203–6.

39. Broad Institute Genome Sequencing Platform & Whole Genome Assembly Team, Jones FC, Grabherr MG, Chan YF, Russell P, Mauceli E, et al. The genomic basis of adaptive evolution in threespine sticklebacks. Nature. 2012 Apr;484(7392):55–61.

40. Kapun M, Fabian DK, Goudet J, Flatt T. Genomic Evidence for Adaptive Inversion Clines in *Drosophila melanogaster*. Mol Biol Evol. 2016 May;33(5):1317–36.

41. Todesco M, Owens GL, Bercovich N, Légaré JS, Soudi S, Burge DO, et al. Massive haplotypes underlie ecotypic differentiation in sunflowers. Nature. 2020 Aug 27;584(7822):602–7.

42. Chen S, Zhou Y, Chen Y, Gu J. fastp: an ultra-fast all-in-one FASTQ preprocessor. Bioinformatics. 2018 Sep 1;34(17):i884–90.

43. Xu H, Luo X, Qian J, Pang X, Song J, Qian G, et al. FastUniq: A Fast De Novo Duplicates Removal Tool for Paired Short Reads. Doucet D, editor. PLoS ONE. 2012 Dec 20;7(12):e52249.

44. Nurk S, Meleshko D, Korobeynikov A, Pevzner PA. metaSPAdes: a new versatile metagenomic assembler. Genome Res. 2017 May;27(5):824–34.

45. Langmead B, Salzberg SL. Fast gapped-read alignment with Bowtie 2. Nat Methods. 2012 Apr;9(4):357–9.

46. Danecek P, Bonfield JK, Liddle J, Marshall J, Ohan V, Pollard MO, et al. Twelve years of SAMtools and BCFtools. GigaScience. 2021 Jan 29;10(2):giab008.

47. Kang DD, Li F, Kirton E, Thomas A, Egan R, An H, et al. MetaBAT 2: an adaptive binning algorithm for robust and efficient genome reconstruction from metagenome assemblies. PeerJ. 2019 Jul 26;7:e7359.

48. Wu YW, Simmons BA, Singer SW. MaxBin 2.0: an automated binning algorithm to recover genomes from multiple metagenomic datasets. Bioinformatics. 2016 Feb 15;32(4):605–7.

49. Alneberg J, Bjarnason BS, De Bruijn I, Schirmer M, Quick J, Ijaz UZ, et al. Binning metagenomic contigs by coverage and composition. Nat Methods. 2014 Nov;11(11):1144–6.

50. Song WZ, Thomas T. Binning_refiner: improving genome bins through the combination of different binning programs. Hancock J, editor. Bioinformatics. 2017 Jun 15;33(12):1873–5.

51. Chklovski A, Parks DH, Woodcroft BJ, Tyson GW. CheckM2: a rapid, scalable and accurate tool for assessing microbial genome quality using machine learning. Nat Methods. 2023 Aug;20(8):1203–12.

52. Uritskiy GV, DiRuggiero J, Taylor J. MetaWRAP—a flexible pipeline for genome-resolved metagenomic data analysis. Microbiome. 2018 Dec;6(1):158.

53. Olm MR, Brown CT, Brooks B, Banfield JF. dRep: a tool for fast and accurate genomic comparisons that enables improved genome recovery from metagenomes through de-replication. ISME J. 2017 Dec 1;11(12):2864–8.

54. Chaumeil PA, Mussig AJ, Hugenholtz P, Parks DH. GTDB-Tk v2: memory friendly classification with the genome taxonomy database. Borgwardt K, editor. Bioinformatics. 2022 Nov 30;38(23):5315–6.

55. Patro R, Duggal G, Love MI, Irizarry RA, Kingsford C. Salmon provides fast and bias-aware quantification of transcript expression. Nat Methods. 2017 Apr;14(4):417–9.

56. Olm MR, Crits-Christoph A, Bouma-Gregson K, Firek BA, Morowitz MJ, Banfield JF. inStrain profiles population microdiversity from metagenomic data and sensitively detects shared microbial strains. Nat Biotechnol. 2021 Jun;39(6):727–36.

57. Grant CE, Bailey TL, Noble WS. FIMO: scanning for occurrences of a given motif. Bioinformatics. 2011 Apr 1;27(7):1017–8.

58. Kılıç S, White ER, Sagitova DM, Cornish JP, Erill I. CollecTF: a database of experimentally validated transcription factor-binding sites in Bacteria. Nucleic Acids Res. 2014 Jan;42(D1):D156– 60.

59. Hyatt D, Chen GL, LoCascio PF, Land ML, Larimer FW, Hauser LJ. Prodigal: prokaryotic gene recognition and translation initiation site identification. BMC Bioinformatics. 2010 Dec;11(1):119.

60. Cantalapiedra CP, Hernández-Plaza A, Letunic I, Bork P, Huerta-Cepas J. eggNOG-mapper v2: Functional Annotation, Orthology Assignments, and Domain Prediction at the Metagenomic Scale. Tamura K, editor. Mol Biol Evol. 2021 Dec 9;38(12):5825–9.

61. Jha N, Kravitz J, West-Roberts J, Camargo A, Roux S, Cornman A, et al. Gaia: A Context-Aware Sequence Search and Discovery Tool for Microbial Proteins [Internet]. 2024 [cited 2025 Jun 4]. Available from: http://biorxiv.org/lookup/doi/10.1101/2024.11.19.624387

62. Weir BS, Cockerham CC. Estimating F-Statistics for the Analysis of Population Structure. Evolution. 1984 Nov;38(6):1358.

63. Nei M, Li WH. Mathematical model for studying genetic variation in terms of restriction endonucleases. Proc Natl Acad Sci. 1979 Oct;76(10):5269–73.

64. Buchfink B, Reuter K, Drost HG. Sensitive protein alignments at tree-of-life scale using DIAMOND. Nat Methods. 2021 Apr;18(4):366–8.

65. Jain C, Rodriguez-R LM, Phillippy AM, Konstantinidis KT, Aluru S. High throughput ANI analysis of 90K prokaryotic genomes reveals clear species boundaries. Nat Commun. 2018 Nov 30;9(1):5114.

66. Price MN, Dehal PS, Arkin AP. FastTree 2 – Approximately Maximum-Likelihood Trees for Large Alignments. Poon AFY, editor. PLoS ONE. 2010 Mar 10;5(3):e9490.

67. Rho M, Tang H, Ye Y. FragGeneScan: predicting genes in short and error-prone reads. Nucleic Acids Res. 2010 Nov;38(20):e191–e191.

68. Eddy SR. Accelerated Profile HMM Searches. Pearson WR, editor. PLoS Comput Biol. 2011 Oct 20;7(10):e1002195.

69. Ondov BD, Treangen TJ, Melsted P, Mallonee AB, Bergman NH, Koren S, et al. Mash: fast genome and metagenome distance estimation using MinHash. Genome Biol. 2016 Dec;17(1):132.

70. Steinegger M, Söding J. MMseqs2 enables sensitive protein sequence searching for the analysis of massive data sets. Nat Biotechnol. 2017 Nov;35(11):1026–8.

71. Matsen FA, Kodner RB, Armbrust EV. pplacer: linear time maximum-likelihood and Bayesian phylogenetic placement of sequences onto a fixed reference tree. BMC Bioinformatics. 2010 Dec;11(1):538.

72. Shaw J, Yu YW. Fast and robust metagenomic sequence comparison through sparse chaining with skani. Nat Methods. 2023 Nov;20(11):1661–5.

